# Simulation-driven discovery of morphology-function relationships in microswimmers

**DOI:** 10.64898/2026.06.01.729272

**Authors:** James F. Cass, Kirsty Y. Wan

## Abstract

For more than a billion years, microorganisms have evolved complex strategies for navigating aquatic habitats, despite the fundamental limitations and constraints imposed by their physical environment. A common theme across these strategies is the use of active slender appendages (cilia, flagella, archaella) to generate self-propulsion. Diverse selection pressures and evolutionary trajectories have driven the emergence of drastically different morphologies of biological microswimmers, each tailored for distinct functions ranging from motility to taxis to prey capture to feeding. Despite the biological and ecological significance of these intricate microscale processes, realistic computational modelling of these organisms and their behaviours is still in its infancy. Here, we present a comprehensive open-source simulation platform for motile microswimmers, that faithfully captures the universal hydrodynamic principles shared by such systems. We illustrate the predictive power and versatility of this approach to resolve and explore morphology-function relationships across different microswimmer species and provide new insights into the diversification of locomotion strategies in early eukaryotes.

## I. INTRODUCTION

The diversity of body plans exhibited by biological microswimmers reflects a myriad of solutions to the unique challenge of life at the microscopic scale [1–3]. In most cases, these organisms actively manipulate their physical body shape to perform a wide range of functions adapted to their ecological niches, guided by very basic nervous systems or even no neurons at all [4–6]. They can navigate complex fluid landscapes, direct their motility in response to various environmental cues (chemical, optical, or mechanical), and identify or capture prey or evade predators [7–13]. These activities are often achieved through the non-reciprocal actuation of multiple slender appendages or protrusions [14–16]. Yet, much of this apparent morphological and functional diversity can be viewed as variations on a common theme from a physical perspective: that is, a single cell or body is equipped with a number of locomotor appendages that are actuated in different ways to accomplish different tasks. Variation comes from morphology (body shape, appendage placement and orientation), and the kinematics of how those appendages beat, coordinate, and interact. This remarkable functional diversity remains relatively underexplored from a quantitative, mechanistic perspective. This makes biological microswimmers rich model systems for understanding how physical structure and motion give rise to adaptive biological function.

How do differences in morphology and beat kinematics across species contribute to differences in motility, feeding, and environmental sensing? Addressing this requires tools that allow systematic exploration of morphology–function relationships across a wide parameter space. While experimental observations have provided a window into these complex behaviours as they unfold in real-time [17–19], and theoretical (or “toy”) models help strip back the inherent complexity of a living organism to clarify the underlying fluid physics [20–23], faithful numerical simulations of microswimmer hydrodynamics offer important complementary insights that bridge the gap between theory and experiment. Simulations can isolate and probe the impact of specific model parameters on the resulting dynamics, perform *in silico* experiments, and make to-scale functional predictions [24–26]. A major challenge is that existing computational approaches can be technically demanding, requiring specialised mathematical knowledge and substantial implementation effort. As a result, exploring new biological hypotheses or morphological variations can be slow and difficult, and inaccessible to researchers without the requisite quantitative training. There is therefore a need for flexible, user-friendly simulation tools that prioritise accessibility and rapid exploration, while retaining sufficient physical realism. Such a framework would accelerate scientific discovery and significantly broaden not only the realm of interesting biological or ecological questions that can be explored, but also extend the range of study species far beyond configurations that are routinely accessible to theory (e.g. pushers, pullers, squirmers) [27, 28].

We structure this paper as follows. First, we introduce `MicroSwimmers.jl`, a free and open-source software package implemented in the Julia language for efficient simulation of microswimmers. This combines high-resolution Stokes flow simulations with a modular, user-friendly interface for designing and testing different microswimmer geometries and appendage shape kinematics. Included are implementations of the swimming problem, predictions of swimming speeds, feeding currents, and particle dispersal, alongside a suite of visualization tools including 3D body locomotion trajectories and flow fields. Next, we demonstrate the versatility of the framework through a set of example research problems in microswimmer hydrodynamics. These examples explore the effects of (i) flagellar beat orientation and its active modulation on 3D swimming trajectories and phototaxis, (ii) geometry and multiciliary coordination when sampling fluid through ciliary bands and (iii) smooth morphological transitions through a biflagellate morphospace. Through these vignettes, we see how a dedicated simulation framework can be used to derive new insights into how organisms exploit fluid-structure interactions for function, with emphasis on how apparently minor changes in behaviour and dynamics can have unexpectedly large effects in the geometrically nonlinear regime.

## II. THE MICROSWIMMERS.JL SIMULATION FRAMEWORK

To support reproducible and extensible exploration of microswimmer dynamics, we developed `MicroSwimmers.jl`, an open-source Julia package for building, simulating, and analysing low-Reynolds-number swimmers (Fig.1(a)). The core of `MicroSwimmers.jl` is a new Julia implementation of the boundary-element regularised-stokeslet method [29, 30]. In this approach, the fluid forces acting on the swimmer surface are represented as a discrete set of regularised point forces (stokeslets) [31, 32], and the resulting linear system relating surface velocities to force distributions is solved (for a detailed description, see Sec. VII). Regularised stokeslets are chosen to prioritise speed and ease of use over extreme accuracy, allowing rapid exploration of morphological and kinematic parameter spaces to understand coupling of microswimmers to their fluid environment and biological functions relative to their niches.

The package extends beyond the core fluid solver to provide a complete simulation pipeline, including tools for swimmer geometry design and high-performance visualisation. A key design principle is modularity. Here, geometries (bodies, grooves, ciliary bands, flagella), beat patterns, and material properties are defined as independent components that can be combined into composite swimmers through a hierarchical design, Fig. 1(b). New flagellar waveform models or body shapes can be added without modifying the core solver, and all components interact through a common low-level interface for geometry access, boundary conditions, and force evaluation. This makes it straightforward to construct new swimmer types, explore morphological parameter spaces and test new biophysical hypotheses.

**FIG. 1.**
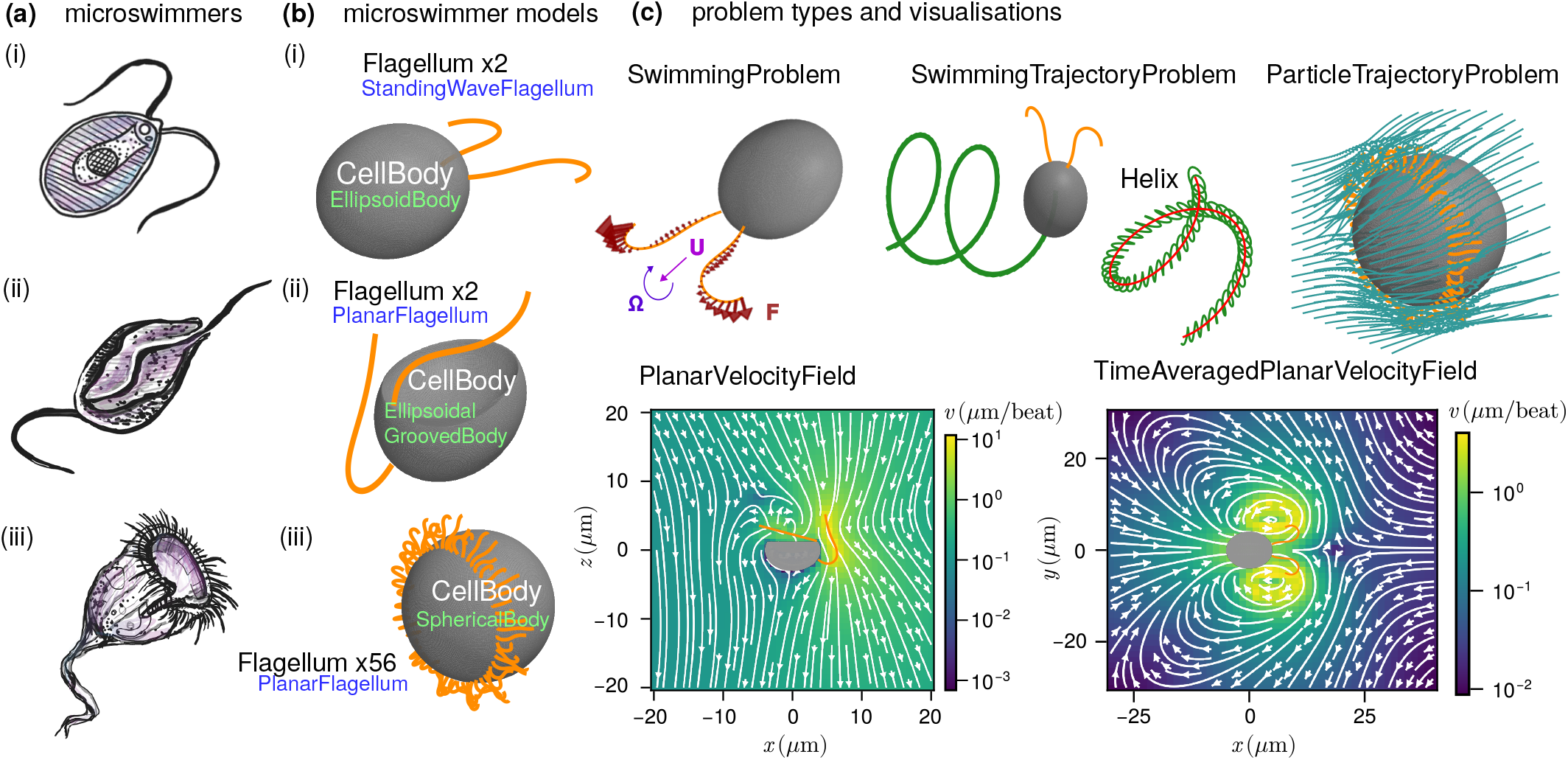
Simulating microswimmer diversity with `MicroSwimmers.jl`. **(a)** Examples of biological microswimmers that vary in body shape, number and arrangement of cilia/flagella and beat kinematics. **(b)** Schematics of the bottom-up hierarchical design process for the microswimmer models used in this paper. Choose a parameterised cell body model (e.g. EllipsoidBody), arrange flagella and choose a kinematic model (e.g. PlanarFlagellum). **(c)** Choose the problem type corresponding to the required boundary conditions e.g. a SwimmingProblem for a free swimmer, and solve numerically (see Sec. VII). Extra functionality is available for analysis including flow field plots, quantifying trajectories through fitted helices and force and torque calculations.

All simulations in this work rely on the same abstract problem interface depicted in Fig. 1(b)-(c). Whether solving instantaneous Stokes problems, time-dependent swimming problems, or calculating derived quantities such as flux, power or torque the user-facing API remains uniform. Time-dependent problems integrate naturally with `DifferentialEquations.jl` for performance and robustness. This unification makes it possible to switch between problem types with minimal code changes and ensures consistent treatment of discretisation, regularisation, and solution across all examples.

We provide a separate package `MicroSwimmersPlots.jl` comprising a suite of visualisation tools (built around `Makie.jl`). These include interactive 3D renderings, animations and flow-field visualisations. Together, these tools enable rapid inspection of simulations, efficient debugging of new models, and clear presentation of results.

## III. RESULTS

In the following section we demonstrate the use of `MicroSwimmers.jl` to derive novel results in three active areas of microswimmer research (i) 3D navigation by beat-plane modulation in *Chlamydomonas* (ii) the effect of geometry and coordination on the filter feeding performance of ciliary bands and (iii) functional impact of slow morphological change on microswimmer behaviour.

### A. Flagellar beat-plane modulation generates zoology of 3D swimming trajectories

Organisms routinely navigate 3D space by dynamically modulating the beat patterns of their swimming appendages [33]. The relationship between appendage actuation and the resulting locomotion behaviour or trajectory is not always straightforward to measure or predict, especially under environmental perturbations [34– 36]. Even for the model biflagellate microalga *Chlamydomonas reinhardtii*, extensively studied in the literature from both experimental and theoretical perspectives, the question of exactly how helical motility trajectories arise from the organism’s coordinated breaststroke gait is only partially resolved (Fig. 2(a)). For example, the apparently planar stroke pattern was revealed to be inherently non-planar using high-speed brightfield [20] and later holographic microscopy [37]. Moreover, a three-bead model where the complex stroke pattern is simplified to beads rotating on circular orbits (each tilted through an angle *β* = 0.3 rad as estimated from experiments), can produce axial rotation of the swimmer along its trajectory [20]. Further asymmetric modulation of the force distribution (represented as a function of phase) reproduced realistic superhelical trajectories characteristic of swimming *Chlamydomonas* when the phase of the flagellar beats was allowed to evolve through interactions with the other parts of the organism.

**FIG. 2.**
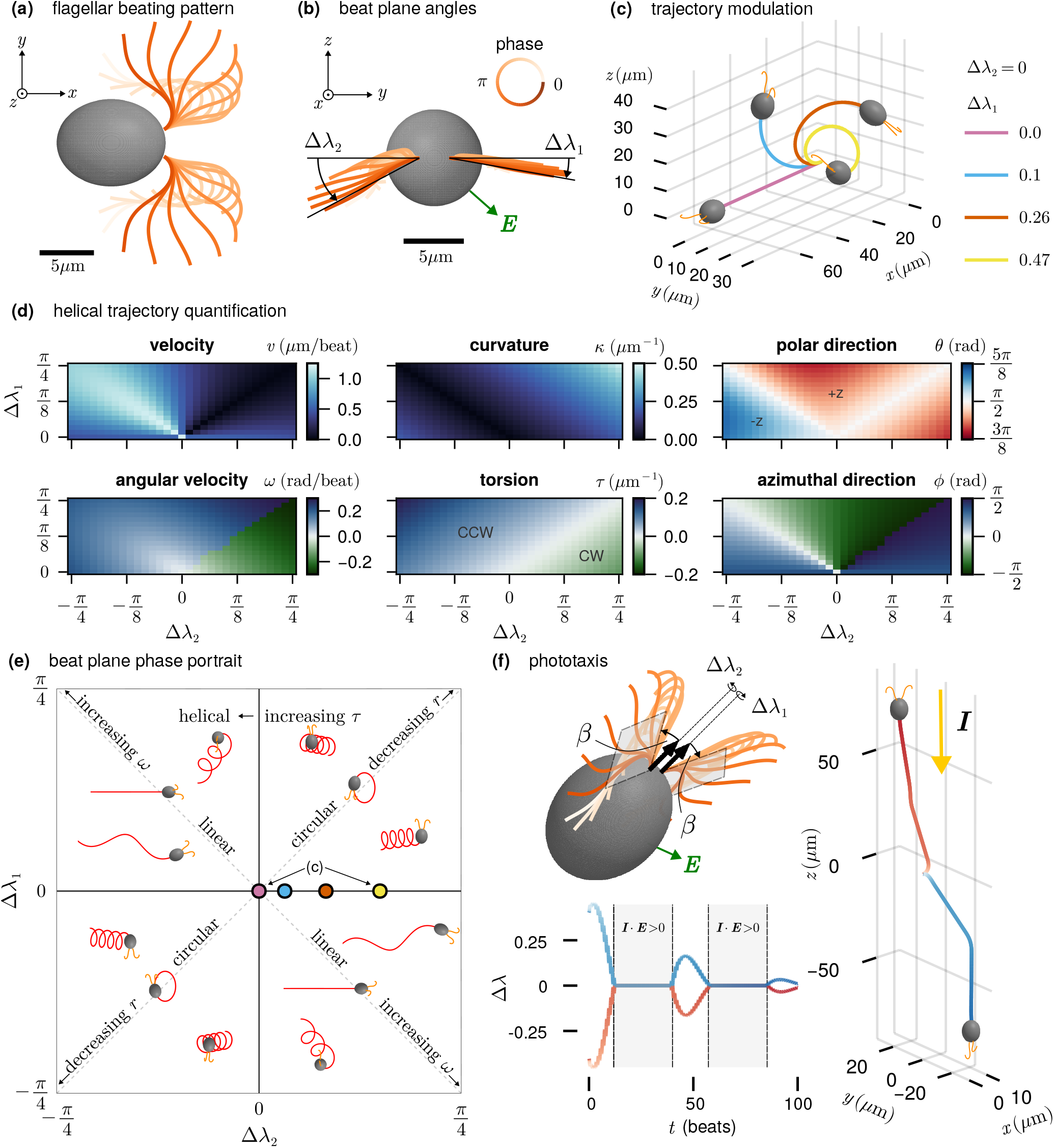
Trajectory space of *Chlamydomonas*. **(a)** `MicroSwimmers.jl` model of *Chlamydomonas* with planar beating pattern (see Sec. VII) **(b)** Beat planes are modulated out-of-plane by rotation angles Δ*λ*_1_ (*cis*) and Δ*λ*_2_ (*trans*) between the beating plane and the *x*-*y* plane (for the *cis* and *trans* flagellum respectively). The eyespot **E** is closer to the *cis* flagellum. **(c)** Example trajectories for single beat plane modulation, transitioning from straight *x*-directed swimming to motion along a helical axis with components in both *y* and *z* directions. **(d)** Fitted helix parameter values and derived quantities for the full (Δ*λ*_1_, Δ*λ*_2_) parameter space. **(e)** Phase portrait representation of trajectories, continued to the lower half plane by symmetry. Markers on the *x*-axis correspond to the simulations in (c). **(f)** Phototactic steering through symmetric beat plane modulation of Δ*λ*_1,2_ around an initial asymmetrically tilted configuration given by angle *β* = 0.3 rad. Note the rotation by *β* is about a distinct axis from the angles Δ*λ*_1,2_. Modulation is due to illumination by light with intensity *I*_0_ = 0.7 (arbitrary units) in the −*z* direction and modulation Δ*λ* ∼ *σ* = ±0.1, where plus/minus response strength *σ* leads to negative/positive phototaxis.

*Chlamydomonas* is an autotrophic green alga that requires light for photosynthesis. Cells perform phototaxis by sensing light intensity gradients temporally (klinotaxis) while navigating a helical path through the surrounding fluid. The physical mechanism of phototaxis has long been thought to depend on asymmetric actuation of the two flagella, termed *cis* and *trans* depending on proximity to the unique eyespot – a rudimentary photoreceptor organelle. Specialised proteins localised to the eyespot region triggers signaling cascades that lead to light-dependent reorientation towards or away from the source of light. Exposure to light has been shown to produce asymmetries in the beat amplitude, frequency and/or phase coordination of the two flagella [38–41]. In the three-bead model, both positive and negative phototactic behaviour can emerge when one or the other flagellum is dominant (produced greater force), suggesting flagellar dominance as a possible control mechanism underlying phototaxis [42]. While force-asymmetry is enough to reproduce quantitatively similar helical trajectories to observed tracks, a recent discovery highlighted the ability for *Chlamydomonas* to modulate the flagellar beat plane itself in a light-intensity dependent manner [43]. While a highly deformable 3D beat pattern has been observed in other flagellates such as *Euglena* [44, 45], beat-plane modulation has not been recognised as a viable control strategy for a cell with bilateral morphological symmetry until now. Here we use `MicroSwimmers.jl` to explore the consequences of beat plane reorientation on three-dimensional swimming trajectories, and ask whether such a strategy, distinct from frequency or amplitude modulation, could still lead to phototaxis.

We begin by specifying a planar beat pattern based on previous waveform tracking data from *Chlamydomonas* [46] (see Sec. VII E). Fig. 2(a) shows the model geometry. We allow the two flagella to execute identical planar beat patterns but differ in the orientations of the beat planes, rotated out of the *x*-*y* plane by angles Δ*λ*_1_ for the *cis*-flagellum and Δ*λ*_2_ for the *trans* (Fig. 2(b)). By varying the two angles, we generate a family of 3D swimming trajectories ranging from straight lines to tight helices. Fig. 2(c) shows the helical trajectories that result from varying only Δ*λ*_1_ while fixing Δ*λ*_2_ = 0. As Δ*λ*_1_ increases, rotating the beat plane further from the *x*-*y* plane, the helices become tighter and the average direction of motion develops positive *y* and *z* components. To systematically explore the space of trajectories that can be reached by varying the two beat plane angles we performed a parameter sweep for Δ*λ*_1_ ∈ [− *π/*4, *π/*4] and Δ*λ*_2_ ∈ [0, *π/*4] (note that the results can be extended to negative Δ*λ*_2_ by symmetry). Trajectories were quantified by fitting to the equation of a helix (see Sec. VII),

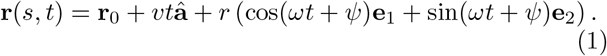

This equation determines the helix (non-uniquely) in terms of 9 parameters (*x*_0_, *y*_0_, *z*_0_, *v, ω, θ, ϕ, r, ψ*). The first three parameters denote a reference point **r**_0_ = (*x*_0_, *y*_0_, *z*_0_) and the directional axis of the helix is given by **â** = (sin(*θ*) cos(*ϕ*), sin(*θ*) sin(*ϕ*), cos(*θ*)). The velocity *v* and angular velocity *ω* are given with respect to this axis direction. We take *v >* 0 so that the swimmer advances in the direction of **â**. With this convention, *ω >* 0 corresponds to right-handed helices and *ω <* 0 to left-handed helices. Fig. 2(d) shows *v, ω, θ* and *ϕ* when varying Δ*λ*_1_ and Δ*λ*_2_. Finally, *r* is the radius of the helix and *ψ* is the phase relative to the two basis vectors **e**_1_ and **e**_2_ (constructed by Gram–Schmidt orthogonalisation) spanning the plane perpendicular to **â**. Derived from these parameters are the curvature *κ* = *r/*(*r*^2^ + *h*^2^) and torsion *τ* = *h/*(*r*^2^ + *h*^2^) where *h* = *v/ω* is the distance moved along the helical axis in a single period (*κ* and *τ* are also shown in Fig. 2(d)). The space of all possible trajectories is summarised in the phase portrait of Fig. 2(e), where combinations of beat plane orientations create straight paths, circles and helices of varying curvature, torsion and handedness, which we describe in detail next (for an example, see Video 1).

The structure of the (Δ*λ*_1_, Δ*λ*_2_) plane is organised by the two diagonal lines of symmetry. Along the line Δ*λ*_1_ = Δ*λ*_2_ the two flagella are rotated symmetrically (either both towards or away from the eyespot), preserving a mirror plane that passes through the cell body between the flagella. Trajectories on this line are circles in the *x*-*z* plane, corresponding to helices with zero *τ*. As |Δ*λ*_1_ | increases along this diagonal in either direction from the origin *κ* increases, producing ever tighter loops. This behaviour matches recent observations of planar circling in *Chlamydomonas* induced by symmetric beat-plane modulation [43].

On the perpendicular diagonal, where Δ*λ*_1_ = − Δ*λ*_2_, the two beat planes are rotated in the same physical direction, preserving a *π*-radian rotational symmetry about the long axis of the body (initially the *x*-axis direction). Trajectories along this line are straight, with the organism rolling around its swimming direction. In helical terms these are curves with zero *κ* and non-zero *τ*. Again the trajectory preserves the unbroken symmetry of the morphology.

The remainder of the parameter plane can be understood in terms of *κ* and *τ* (Fig. 2**b**). As we have seen *τ* vanishes (planar circles) on the line Δ*λ*_1_ = Δ*λ*_2_; lines parallel to this diagonal are lines of constant nonzero *τ* whose magnitude increases with distance from the diagonal. The sign of *τ* gives the intrinsic handedness of the helix around its axis. Conversely, trajectories on the diagonal Δ*λ*_1_ = − Δ*λ*_2_ have zero *κ* and parallel lines are constant *κ* helices, with *κ* increasing with distance from the diagonal on either side.

In Fig. 2(c) we saw that modulating a single beat plane already produces trajectories with components in all three spatial directions. In the full parameter space a single beat plane is varied on the coordinate axes of the (Δ*λ*_1_, Δ*λ*_2_) plane. These points show the largest deviations of the polar angle *θ* of the axis vector **â** from *θ* = *π/*2 (the *x*-*y* plane) corresponding to the strongest excursions of the trajectories in the *z* direction. The azimuthal angle varies between *ϕ* = − *π/*2 to *ϕ* = *π/*2 corresponding to motion that steers the swimmer in the *x*-*y* half plane of positive *x* values. The change in handedness across Δ*λ*_1_ = Δ*λ*_2_ already discussed manifests here as a discontinuity in the azimuthal angle, where trajectories on either side of the discontinuity progress in opposing directions.

We next test whether temporal modulation of the beat angles alone suffices to generate phototactic steering. We applied a light-dependent modulation rule on top of an initial configuration inspired by evidence that the intrinsic beat plane of *Chlamydomonas* is itself non-planar (prior to any modulation) [20]. Fig. 2(f) shows this initial configuration, where the beat planes are tilted by an angle *β* = 0.3 rad away from the *x*-*y* plane, guaranteeing body rotation around the swimming direction that leads to periodic illumination of the eyespot [20]. The angle *β* differs from Δ*λ*_1,2_ in the axis about which the beat plane is rotated. The modulation is set by the intensity of light *S*(*t*) detected at the eyespot, produced by a constant light stimulus **I** = − *I*_0_**e**_*z*_ (yellow vector). The eyespot location, closer to the *cis* flagellum, is shown by the vector **E** in Fig. 2(b) and (f). The intensity at the eyespot is

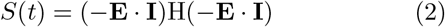

which is only non-zero when the eyespot is exposed to the light (H is the Heaviside function). When the eyespot enters the illuminated region, the beat planes are modulated symmetrically to Δ*λ*_1,2_ = Δ*λ* = *σ* log(1 + *S*(*t*)), where *σ* is the strength of the response. Fig. 2(f) shows the trajectories and time dependence of Δ*λ* for both a positive and negative response, *σ* = 0.1 or *σ* = − 0.1. For positive Δ*λ* both flagella move towards the eyespot. This modulation was observed experimentally under high light intensity [43]. The effect is to reorient the helical axis away from the light source (negative phototaxis, blue lines). The reorientation happens within intervals of time where the eyespot is illuminated, corresponding to **E** · **I** *<* 0. Conversely, with beat plane modulation away from the eyespot (corresponding to *σ <* 0 and observed under low light intensity conditions) the trajectory is positively phototactic, with the trajectory simply mirrored through the *x*-*y* plane.

Together, these results show that modulating the two beat plane angles provides direct control over the orientation of the swimming axis. Further, (symmetric) beat plane modulation even in the absence of force asymmetry is a viable and effective strategy for phototactic reorientation for a *Chlamydomonas*-like microswimmer.

### B. Morphology and ciliary dynamics govern filter feeding performance

Many planktonic species, ranging from ciliates to animal larvae, rely on flows driven by the collective dynamics of cilia to create currents for filter feeding [21, 47]. The ability to acquire nutrients by feeding or engulfing other organisms and particulates is considered to be a critical cellular innovation, that enabled single-celled eukaryotes to attain larger size and greater behavioural complexity [6]. Distinct ciliary architectures and feeding apparatuses have evolved to fulfil this flow-generation function, which is especially important in larger organisms to overcome diffusive bottlenecks associated with passive nutrient uptake [48]. Different considerations apply when considering the energetics of swimming versus feeding in motile or sessile organisms, which exhibit different configurations and body coverage of cilia.

A recurring motif is the ciliary band – dense rings of cilia, often found adjacent to a mouth or pharyngeal opening, that beat to drive flows of nutrients or prey towards the cell. The dependence of feeding performance on body morphology, band placement and ciliary coordination remains incompletely characterised. Previous modelling studies of suspension feeding in ciliates have mainly considered the spherical squirmer model, prescribing a tangential slip velocity at the body surface as a simplified representation of the induced ciliary flows [21]. Numerical simulations involving more complex geometries and filament dynamics would help clarify how ciliary arrangement and orientation shape these flow patterns.

Here, we study two idealised filter-feeder designs modelled on sessile organisms (body-tethered), one with a spherical body and a single band of cilia, and a second morphology where the region inside the ciliary band (a spherical cap) is inverted to form a cavity of the same surface area, while keeping everything else the same (Fig. 3(a)). The second case more closely resembles real filter feeders, and is intractable analytically within the spherical squirmer framework. Both swimmers have a radius of *a* = 20 μm, and a ciliary band at height *h >* 0 (where *h* = 0 corresponds to a band on the equator; the band approaches the anterior pole as *h/a* approaches unity) with radius 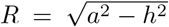. Power strokes are directed along the positive *x* axis. We fix the spacing between cilia along the band to be 2 μm, meaning that ciliary bands with larger radii contain more cilia. The flagellar beat dynamics are fixed (see inset), while the coordination pattern of the cilia within the array is allowed to vary.

**FIG. 3.**
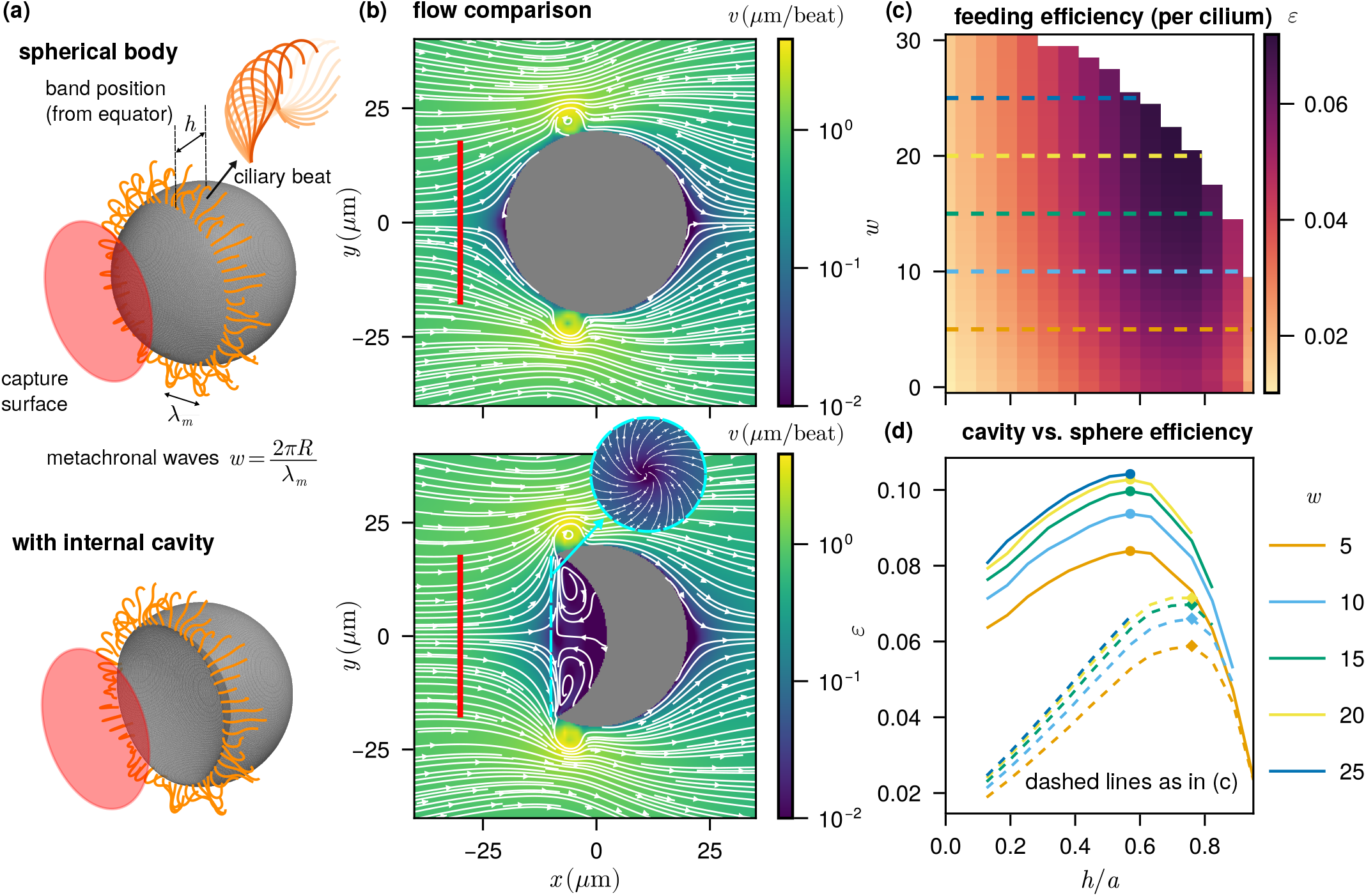
Filter feeding through a ciliary band. **(a)** (i) (top) A spherical filter feeder model with cilia spaced 2 μm apart on a ciliary band. We calculate the feeding flux *Q* through the capture disk surface shown. We vary the height of the band *h* from the equator and the number of metachronal waves *w* along the band. (bottom) Second model with an indented cavity region forming a mouth of the same surface area as the spherical cap in the base model. **(b)** Planar flow structures for the two body types, averaged over a flagellar beat cycle. **(c)** The efficiency of feeding, given by *ε* = *µQ*^2^*/P*, through the capture surface as a function of band height *h* and metachronal waves *w*. Dashed lines are marked for comparison with panel (d). **(d)** Improved feeding efficiency for the body with internal cavity (solid lines) relative to the spherical case (dashed lines).

We vary the band height *h* and the number of metachronal waves *w* = 2*πR/λ*_*m*_ along the band, where *λ*_*m*_ is the metachronal wavelength (Fig. 3(a)), quantifying performance by the volume flux (clearance rate)

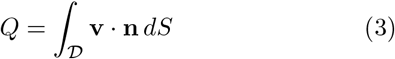

through a capture disk 𝒟 with the band radius *R* positioned 10 μm from the anterior pole. Following Osterman and Vilfan [49] we measure feeding efficiency per cilium by 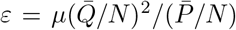, where the over bar denotes a time average, *N* is the number of cilia in the band and the instantaneous hydrodynamic power is

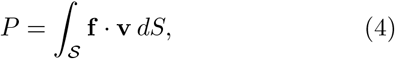

integrated over the swimmer surface 𝒮 (see Methods for the numerical calculation of integrals). This metric accounts for the power required to generate a given flux and avoids the confounding effect of increasing cilia number with band circumference. Note that *ε* as defined here has the units of an effective “feeding volume” which could be made dimensionless through dividing by *a*^3^ (this would only rescale the results presented).

The time-averaged flow structures for the two body types (Fig. 3(b)) reveal a qualitative difference introduced by the cavity. In both cases, representative fluid particle trajectories show fluid drawn toward the mouth and swept across the ciliary band (also see Fig. 1(c) and Video 2). For the spherical body, incoming fluid must pass through a high-drag region near the surface before reaching the ciliary band. For the body with an internal cavity, in contrast, fluid approaching the mouth encounters a stagnation point at the cavity entrance, with flow on either side directed toward the ciliary band. Fluid within the cavity circulates in a vortex ring (seen by rotating the lower flow structure in Fig. 3(b) in three dimensions), generating an effective slip surface at the mouth entrance that assists transport towards the band. Since there is no mass flux through the cavity walls, only fresh exterior fluid is drawn toward the feeding band.

The efficiency *ε* as a function of band height *h* and metachronal wavenumber *w* is shown in Fig. 3(c,d). For the spherical body, efficiency attains a clear maximum near *h* = 14 μm, reflecting a balance between two competing effects: bands near the anterior pole subtend a small capture area, reducing throughput, while bands near the equator force incoming fluid to traverse a larger low-velocity region before reaching the cilia (see Video 2). The cavity geometry improves the feeding efficiency dramatically (by close to a factor of two), while shifting the optimal band height away from the pole and closer to the equator. Increasing the number of metachronal waves *w* enhances efficiency monotonically across all band heights (Fig. 3(c)), consistent with findings that coordinated phase differences between neighbouring cilia improve fluid transport [25, 50]. This effect diminishes, however, as *w* approaches the maximum number of metachronal waves that can be supported by the number of cilia on the band.

The combined dependence on band height and metachronal wavenumber highlights that both the geometry of the ciliary band and the spatiotemporal coordination of its beating affect overall feeding performance. The results also suggest that the evolution of invaginations or inverted curvature regions associated with ciliary bands, a morphological motif also observed at the scale of colonial choanoflagellates [51], may serve to increase fluid and nutrient uptake. In other organisms, particularly trochophore larvae of marine invertebrates, where ciliary bands are used for propulsion instead of feeding [52], the ciliary band may still be positioned equatorially.

### C. Smooth morphological and kinematic changes drive major swimming transitions

In this final example we consider the effect of smooth morphological variation on swimming behaviour across biflagellate protists. The biflagellate condition, namely, when a cell or organism has precisely two flagella that may serve either identical or distinct functions, is highly prevalent across the eukaryotic tree of life [27, 53]. Recent phylogenetic evidence suggests that the last eukaryotic common ancestor (LECA) may have possessed a biflagellate morphology closely resembling modern-day excavates – a paraphyletic group of early-branching unicellular eukaryotes that may have retained certain ancestral traits [54]. This includes presence of a (sometimes vaned) flagellum that beats vigorously inside a feeding groove [55]. Most excavates are biflagellated, with two heterodynamic flagella that are markedly asymmetric in morphology, function as well as behaviour. They typically do not synchronise their movements and beat at well separated frequencies [55]. In contrast, the late-diverging Chlorophyte alga *Chlamydomonas* exhibits a well-characterised synchronized breaststroke with a pair of symmetric flagella of equal length (see also Sec. III A above). This implies substantial changes in flagellar arrangement and body morphology must have occured during divergence of ancestral biflagellates into modern-day *Chlamydomonas*.

We construct an idealised biflagellate model parameterised in such a way as to enable smooth variation between an excavate-like morphology (asymmetric flagella with groove) and a *Chlamydomonas*-like morphology (symmetric flagella, no groove), not intended as a faithful reconstruction of this evolutionary transition, but as a controlled axis of morphological variation (Fig. 4(a)). The body is a fixed prolate spheroid with semi-axes *a* = 3.9 μm and *b* = *c* = 2.2 μm, with flagella of fixed length 11 μm. The feeding groove is constructed by subtracting an ellipsoid with identical dimensions from the body ellipsoid, that is shifted in the positive *z* direction a distance *d* relative to the body. Five parameters are linearly interpolated between the two endpoint morphologies: the aforementioned groove ellipsoid offset *d* (from 2.6 μm to 6.1 μm), the azimuthal attachment angle *β* of the flagella relative to the anterior pole (moving through *π* radians), the average (static) dimensionless curvature *C* of the posterior flagellum (shown by dotted lines), and finally the two beat plane angles *R*_*y*_ and *R*_*x*_ moving through angles *π* and *π/*2 respectively. The two flagella are assumed to share the same beat pattern throughout with a fixed separation between their attachment points. A larger spacing than the adjacent basal bodies of extant excavates is used, as this more naturally recovers the *C. reinhartii* -like morphology without introducing further parameters. We then define a single parameter *α* ∈ [0, 1] which controls the simultaneous linear interpolation of the set of parameters {*β, d, R*_*x*_, *R*_*y*_, *C*}. This parameterisation allows gradual changes in morphology and beat geometry to be explored in a controlled way.

**FIG. 4.**
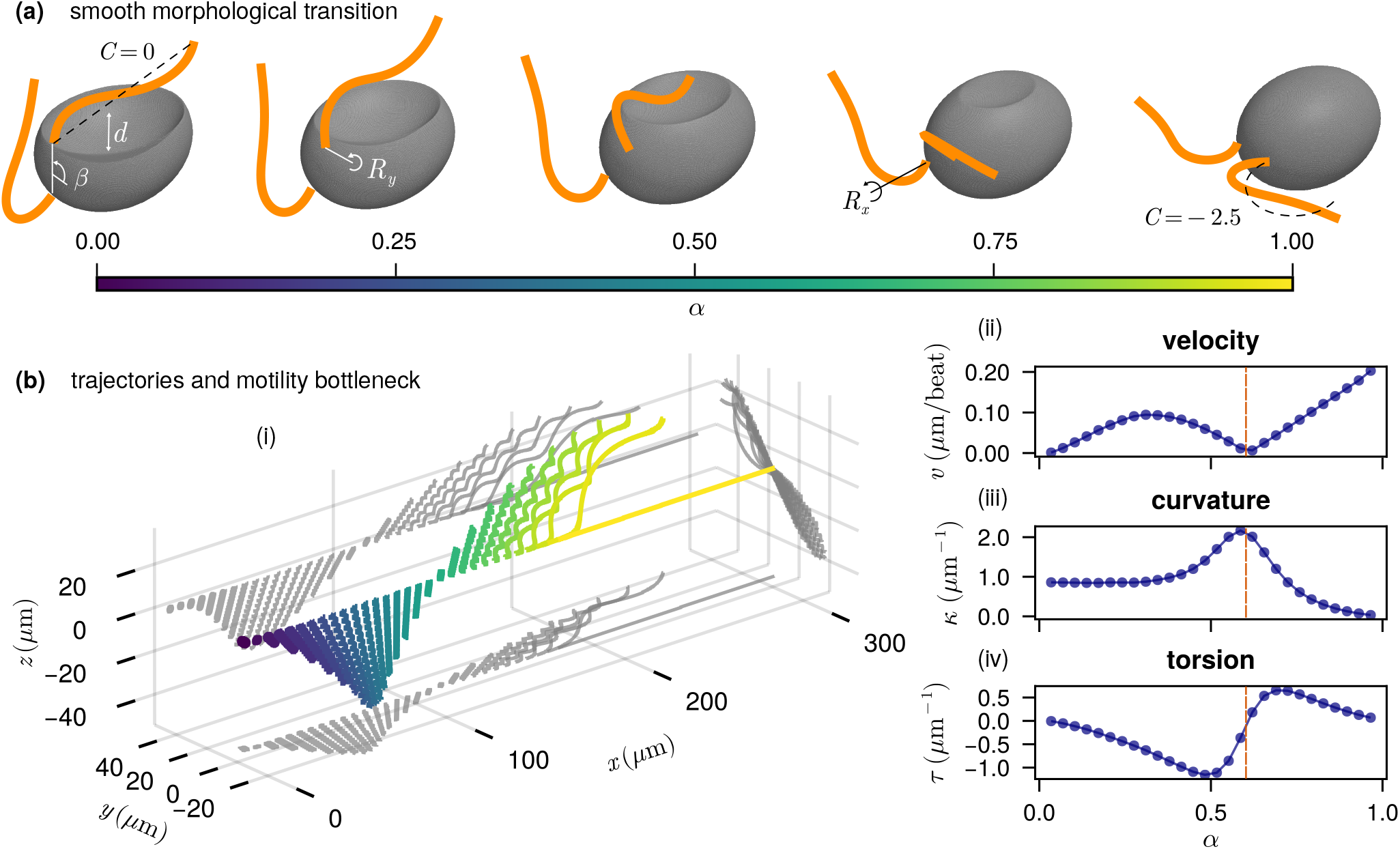
Smooth transition from a typical excavate like morphology to *Chlamydomonas*. **(a)** Shapes are parameterised by 5 parameters (*β, d, R*_*x*_, *R*_*y*_ and *C*). The colour bar shows *α* ∈ [0, 1] representing a linear interpolation of the five parameters taking one morphology to the other. **(b)** Helical trajectories over 500 beat cycles as *α* varies, (starting points spaced out along *x* for visualisation). **(c)** Quantification of the helical trajectories fitted to trajectories in (b), highlighting a motility breakdown point where *v* = 0.

Swimming trajectories along this axis of variation reveal that gradual morphological changes can produce dramatic and non-monotonic shifts in behaviour (Fig. 4(b)). The endpoint morphologies have degenerate trajectories reflecting their symmetries: circles for the excavate-like configuration and a straight line for the *Chlamydomonas*-like breaststroke. Along the interpolation path the helix curvature *κ* intially remains constant before undergoing a sharp increase (Fig. 4(b)(iii)), at which point motility breaks down (*v* = 0, Fig. 4(b)(ii), dashed line). Fig. 4(b)(i) shows trajectories for *α* below this critical value make progress in the negative *y*, negative *z* directions, whereas for *α* higher than this value there is progress in the positive *x, y* and *z* directions. This swimming direction reversal is accompanied by a chirality switch captured by a sign change in torsion *τ* (Fig. 4(b)(iv)). As the *Chlamydomonas*-like gait is approached the curvature decreases until only *x* motion remains.

These results highlight that seemingly smooth morphological changes can lead to abrupt and non-monotonic changes in swimming behaviour. Persistent forward motion versus recurrent high-curvature paths may confer different advantages for efficient phototaxis or for foraging near surfaces respectively. Our findings therefore suggest that simple monotonic increases in adaptive performance are unlikely: transitions between distinct flagellar organisations may have required navigating behavioural bottlenecks along the way. Intermediate states do not resemble any extant protist species, perhaps due to the extremely low motility performance.

## IV. DISCUSSION

We developed the `MicroSwimmers.jl` framework, and used it to explore three representative problems: the effect of beat-plane modulation on the swimming trajectories and phototaxis of *Chlamydomonas*, the influence of ciliary-band morphology and coordination on filter feeding, and the functional consequences of smooth morphological variation on biflagellate swimming motility. Across these systems, the mapping from geometry and coordination to swimming or feeding performance is highly nonlinear, with sharp transitions and strong symmetry constraints shaping the accessible behavioural space. This demonstrates how small morphological or kinematic modifications can produce disproportionately large changes in the function and behaviour of microswimmers.

For *Chlamydomonas*, phototactic reorientation is typically attributed to differences in the force generated by the *cis* and *trans* flagella, resulting from asymmetric beat amplitudes (and possibly also frequency and beat extension) depending on the direction of light stimulus [38, 39, 42, 56]. Our results show a new strategy in which beat-plane modulation alone is sufficient to generate direction-modulated trajectories. By tilting the effective beating planes in response to light intensity perceived at the eyespot, the swimmer can reorient its helical axis towards or away from a stimulus without invoking force asymmetries. The base values were motivated by observations in experiments with tethered cells [20, 37], and our modulation rule gives rise to similar magnitudes as those observed due to light intensity changes [43]. From the perspective of cell physiology, how is beat-plane modulation implemented, and how does it interact with force dominance or phase asymmetry *in vivo*? Whether the organism exploits this mechanism in addition to force dominance remains an open question, along with the origins of the asymmetry between *cis* and *trans* flagella. Our results highlight the need for targeted experiments capable of separating these modes of control, with sufficient resolution to measure non-planar flagellar dynamics in freely swimming cells in response to controlled photo-stimulation.

The filter-feeding example demonstrates how organism morphology, ciliary-band placement and coordination jointly determine clearance rates. Although the volume flux increases with band circumference, this effect is counteracted by the formation of a large low-velocity region as the band approaches the equator, increasing the distance between the band and the feeding surface. As a result, feeding efficiency exhibits a clear optimum at a band height midway between the anterior pole and equator, consistent with predictions using a banded squirmer model [57]. Simulations allowed us to go beyond the spherical geometry to study the effect of invagination, with a striking feeding performance gain resulting from a qualitatively different flow structure. Metachronal coordination enhances performance across all band positions, consistent with theoretical predictions that coordinated phase gradients can improve transport efficiency. These results reinforce the idea that both geometry and coordination must be considered together when assessing the cilia-driven feeding strategies of sessile and motile organisms.

Finally, our excavate–*Chlamydomonas* transition highlights how gradual morphological change can lead to abrupt behavioural shifts. Although the interpolation between flagellar attachment angle, groove depth, beat-plane parameters, and posterior-flagellum curvature is smooth, the resulting swimming trajectories display a curvature spike indicating a breakdown in motility. These behavioural bottlenecks suggest that not all morphological trajectories through shape space are equally viable: evolution may have been constrained to paths that avoid regions where propulsion becomes inefficient or unstable, or else opted for abrupt rather the smooth transitions in cellular ultrastructure. This underscores the value of coupling motility simulations with phylogenetic and morphological data, as behavioural constraints may help to narrow down plausible evolutionary scenarios. Indeed, the behavioural bottleneck we identified likely corresponds to a fitness valley, consistent with the theory that *Chlamydomonas*, a highly-derived green algal lineage, underwent radical cytoskeletal reorganization particularly of its flagellar apparatus [58, 59], when it traded off the phagotrophic lifestyle of its ancestors for efficient swimming and photosynthesis.

By combining high-resolution Stokes flow simulations with a flexible computational framework, we have shown how subtle changes in beat geometry, ciliary coordination, or organismal shape can lead to qualitatively different swimming and feeding behaviours. These results highlight the value of computational tools that bridge detailed biophysics with ecological and evolutionary questions, and suggest that the organisation of flagella and cilia may be shaped as much by behavioural constraints as by morphology or genetics. The `MicroSwimmers.jl` framework unlocks new discoveries and enables new biological hypotheses to be tested by providing a modular, extensible environment in which geometries and beat patterns can be combined with consistent numerical solvers and analysis tools. Future directions include inference of beat parameters from experimental trajectories, integration with self-organised models of ciliary axonemal beat dynamics [60, 61], and the systematic exploration of morphology–behaviour maps across broader phylogenetic groups. Many extensions to the basic framework are in progress or planned e.g. interactions between multiple microswimmmers or colonies and custom boundary geometries. We hope that the user-friendly approach we have prioritised will lead to community uptake and participation in the project through github.com/micromotility-lab/MicroSwimmers.jl.

## Supporting information

Movie 2: Filter-feeding by a spherical microswimmer (tethered) in which metachronal waves of cilia drive fluid particles across the ciliary band.

Movie 1: Simulation of a Chlamydomonas cell swimming on a helical trajectory under beat plane modulation.

## V. ACKNOWLEDGMENTS

We would like to thank Alastair Simpson, Thomas Kiørboe, Sei Suzuki-Tellier and members of the Wan lab for stimulating discussions. This work was funded by the European Research Council (ERC) under the European Union’s Horizon 2020 research and innovation programme grant 853560 Evomotion and the Human Frontier Science Program RGP014/2024 (KYW). The authors would like to acknowledge the use of the University of Exeter’s Advanced Research Computing facilities in carrying out this work.

## VI. DATA AND CODE AVAILABILITY

The `MicroSwimmers.jl` package, analysis scripts, and data required to reproduce the results in this paper are archived at https://doi.org/10.5281/zenodo.20340772 [62]. The live package repositories are hosted at github.com/micromotility-lab/MicroSwimmers.jl

## VII. METHODS

### A. Boundary element method with regularised stokeslets

The boundary element method is based on the reciprocal theorem of Stokes flow. Assume the pairs (**u, *σ***) and (**u**^′^, ***σ***^′^) represent the velocity and stress tensors of physical and auxiliary flows respectively, that are solutions of the Stokes equations

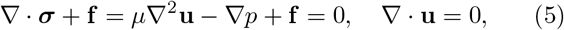

where *p* is the pressure, *µ* the dynamic viscosity and **f** is a body force per unit volume. The product rule (along with the symmetry of the stress tensor and the incompressibility of the fluids) leads to the identity [63] (switching to index notation)

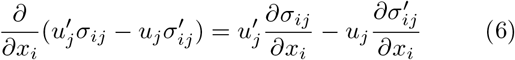

For the physical flow with no body forces (**f** = 0), we have *∂σ*_*ij*_*/∂x*_*i*_ = 0 from Eq.(5). The boundary integral equation can then be derived by taking the auxiliary flow solution as the *stokeslet* (the solution to a dirac delta body forcing **f** = *δ*(**x** − **y**)**F**) and using the divergence theorem to integrate over the volume.

In contrast to the exact Greens function approach, regularised stokeslets are solutions to the Stokes equations forced by a smoothed point force of width *ϵ* applied at **y** in an otherwise unbounded domain [31, 32]. A commonly used regularised stokeslet solution has velocity response 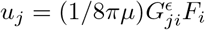 where the regularised stokeslet tensor is given by

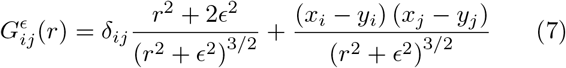

with *r* = |**x** − **y**|. This velocity field in this case satisfies

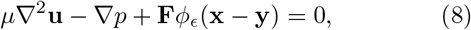

for an arbitrary force **F** (that will cancel below) with the regularised forcing

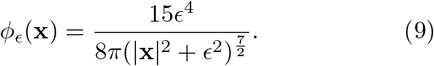

Using the regularised stokeslet solution for (**u**^′^, ***σ***^′^) in the reciprocal theorem (equation (6)), integrating over the (**u, *σ***) problem domain and applying the divergence theorem yields the approximate boundary integral equation

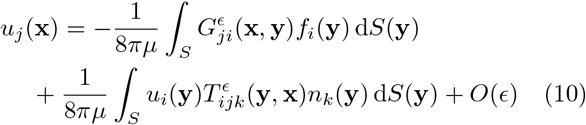

where **f** = ***σ*** · **n** is the boundary traction. The first term on the right-hand side is the single-layer potential and the second is the double-layer potential. As is common in regularised-stokeslet BEM, we neglect the double-layer contribution, obtaining

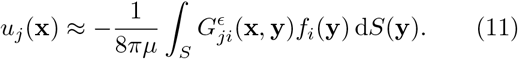

For a prescribed flagellum velocity this has been shown to be a good approximation [64]. Solving this linear equation for a prescribed velocity **u**(**x**) on the boundary gives the unknown traction **f** (**x**), with which the fluid velocity at any point in the domain can be calculated using equation (11).

### B. Nearest-neighbour discretisation

To evaluate the boundary integral equation (11) numerically we use the nearest-neighbour discretisation scheme introduced by Smith *et al*. [29, 30]. The key idea is that the force distribution typically varies on a longer lengthscale than the rapid variation of the regularised stokeslet kernel. We therefore discretise the surface or filament (in the case of flagella) using two sets of points: a fine set of *quadrature points* to capture the rapid variation and a coarser set of *force points*. Each quadrature point *q* is associated with exactly one force point *n*—its nearest neighbour in Euclidean distance (in a reference configuration), via the mapping

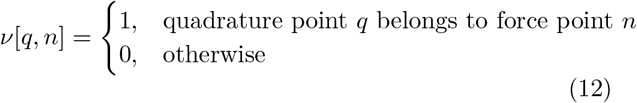

The boundary traction is assumed constant over the neighbourhood of a force point, and we further assume that the area or line element associated with a quadrature point varies slowly within the force point neighbourhood. It is important to treat the parts of a microswimmer separately (body, groove, flagellum etc.) when assigning nearest neighbours, so that no patch spans a region where the surface metric changes rapidly. The discretised version of equation (11) is

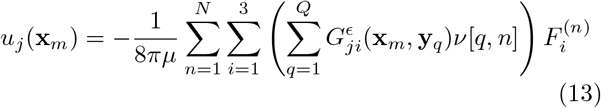

where *m* indexes force points. The unknowns 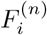 are given by

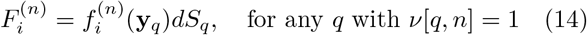

where 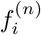 is the (assumed constant) traction at force point *n* and *dS*_*q*_ is the area element at quadrature point *q*, assumed uniform within the patch. Since the unknowns are 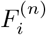, the linear system has 3*N* degrees of freedom regardless of *Q*. Note that the quadrature weights are absorbed into the unknowns, so they do not appear explicitly in the discretised system.

### C. Other implementations of the boundary-element regularised-stokeslet method with nearest-neighbour discretisation

The method originates with a MATLAB implementation due to Smith [29]. Follow-up papers extended the method to add swimming problems [30], the double-layer potential [64] and Richardson-extrapolation enhanced numerics [65]. More recently the same group published an implementation using C++ that addresses some of the issues with speed and memory usage in the MATLAB version [66]. Our Julia implementation similarly addresses the main issue of forming an intermediate matrix requiring large memory usage. The MATLAB-style syntax of Julia and the design and visualisation infrastructure we provide around the simulation pipeline should make this powerful method more accessible to those without C++ programming experience.

### D. Computing forces, torques and velocity fields

Once the force-node unknowns **F**^(*n*)^ are obtained, the total hydrodynamic force and torque on any part of the swimmer are reconstructed by summing the contributions of the quadrature nodes associated with each force node. If the nearest-neighbour mapping assigns quadrature point *q* to the force node *n*, the total force is

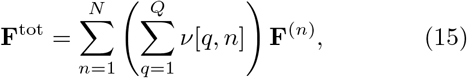

and the total torque about a reference point **x**_0_ is

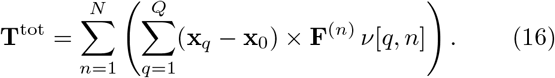

To evaluate the fluid velocity at a point **y** in the domain, we reuse the single-layer representation, equation (13)

### E. Implementing a flagellar beat pattern

A flagellum in `MicroSwimmers.jl` is modelled as a parameterised, time- and arclength-dependent shape, specified through its tangent angle *θ*(*s, t*) along the flagellum. Here *s* ∈ [0, 1] is a dimensionless arclength coordinate along a flagellum of length *L*, so that the position of the flagellum in its local frame is obtained by integrating

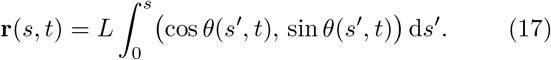

Two slightly different beat-pattern models are used in this paper: a `PlanarFlagellum` and a `StandingWaveFlagellum`.

The `PlanarFlagellum` model represents the tangent angle as a steady intrinsic curvature plus a travelling-wave deformation with a spatially modulated amplitude [30]

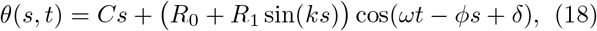

where *C* sets the static bend, *ω* is the beat frequency, *ϕ* controls the spatial phase gradient along the flagellum (and hence the wave propagation direction and wavelength), and *δ* is a phase offset used to coordinate the relative phase of multiple flagella as in Sec. III B. The amplitude envelope *R*_0_ + *R*_1_ sin(*ks*) allows the beat amplitude to vary along the flagellum: *R*_0_ sets a uniform amplitude, while *R*_1_ and *k* together control its spatial modulation. This form provides a compact representation of travelling-wave beats such as those used by sperm and many uniflagellate swimmers when *C* = 0 and a power stroke/recovery stroke *Chlamydomonas*-like beat when *C* ≠ 0 [67]. The examples in Sec. III B and Sec. III C use a `PlanarFlagellum` model with the parameters *L* = 11 μm, *C* = − 2.46 rad, *R*_0_ = 0.35 rad, *R*_1_ = 1.5 rad, *k* = 0.7, *ϕ* = 6.22, and *ω* = 2*π* so that one simulation time unit corresponds to the beat period.

The `StandingWaveFlagellum` model represents the tangent angle as a steady intrinsic curvature plus standing-wave deformations [68],

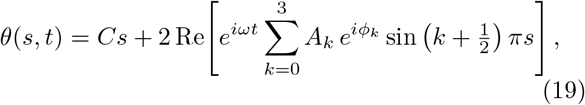

where *C* sets the static bend, *ω* is the beat frequency, and {*A*_*k*_, *ϕ*_*k*_} are the amplitudes and phases of the standing-wave modes. The spatial basis sin 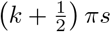 enforces zero tangent angle at the base (*s* = 0) and a free end at the tip (*s* = 1). For the *Chlamydomonas* simulations in Sec. III A the amplitudes and phase were obtained from a discrete Fourier decomposition of experimental beat data [46]. The parameters used were *L* = 11 μm, *C* = − 2.20, and standing-wave amplitudes and phases (*A*_0_, *ϕ*_0_) = (0.598, 0.537), (*A*_1_, *ϕ*_1_) = (0.566, 0.158), (*A*_2_, *ϕ*_2_) = (0.158, 2.197), again with frequency *ω* = 2*π*.

Once a beat-pattern model is defined, a complete flagellum is constructed by attaching it to a cell body at a specified location and orientation. For example, the *Chlamydomonas* model has two flagella placed symmetrically about the anterior pole of the ellipsoidal body at positions (*a* cos *α*, ± *b* sin *α*, 0), with their beat planes initially parallel to the *x*–*y* plane and rotated outward by an angle *γ* = 0.5 rad about the *z*-axis so that the flagella emanate perpendicular to the cell body (Fig. 2(a,b)). Coded examples for constructing flagella, complete swimmers and solving the various problem types are provided in the Zenodo archive [62].

### F. Flux and power integrals

We quantify feeding by computing the volumetric flux through a circular feeding disk of radius *R* centred at (*x, y*_0_, *z*_0_), with normal aligned with the *x*–axis (Fig. 3(a)). The instantaneous flux is

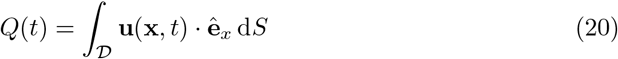

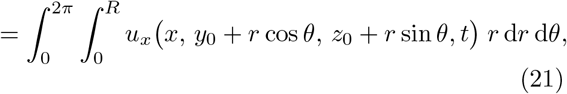

where *u*_*x*_ is the *x*–component of the fluid velocity and *R* is the disk area. Numerically, we evaluate (21) using tensor-product Gauss–Legendre quadrature in *r* and *θ*, with the factor *r* accounting for the polar area element. *Q*(*t*) is then averaged across a beat period.

The total hydrodynamic power expended by the swimmer at time *t* is

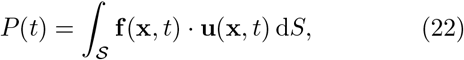

where **f** is the surface traction and 𝒮 denotes the swimmer surface. Using the nearest-neighbour discretisation described above, this integral is approximated by a sum over quadrature points,

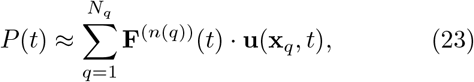

where **x**_*q*_ are the quadrature points, *n*(*q*) is the index of the force point associated with **x**_*q*_ by the nearest-neighbour mapping, and **F**^(*n*)^ is the corresponding force contribution (which already incorporates the quadrature weight).

### G. Fitting the parameters of a helix to a simulated trajectory

We fit the simulated swimming trajectories to a helical curve of the form Eq. (1). To obtain the parameters of the helix from a simulated trajectory, we first construct an initial guess using a principal–component analysis and then refine this using nonlinear least-squares, as follows. Given a trajectory {**r**(*t*_*k*_)}, we subtract the mean position and form the data matrix with columns 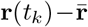. We then compute its singular value decomposition and take the first left singular vector **â** as an estimate of the helical axis direction. An orthonormal transverse basis {**e**_1_, **e**_2_} spanning the plane perpendicular to **â** is obtained by projecting a fixed Cartesian basis vector onto this plane and normalising, then setting **e**_2_ = **â** × **e**_1_.

The trajectory is then expressed in this moving frame. Writing

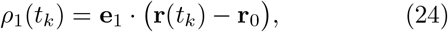

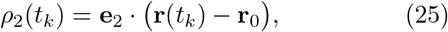

we define the phase *ϕ*(*t*_*k*_) = atan2(*ρ*_2_(*t*_*k*_), *ρ*_1_(*t*_*k*_)), which is approximately linear in time. The initial point **r**_0_ is estimated by projecting the centroid 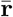 onto the plane perpendicular to **â**, giving 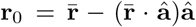. A linear regression *ϕ*(*t*) ≈ *ωt* + *ψ* provides initial estimates for the angular frequency *ω* and phase offset *ψ*. The radius is estimated from the spread of the trajectory in the transverse directions, and the axial speed *v >* 0 from the displacement along **â** over the duration of the trajectory.

These initial parameters are used as a starting point for a nonlinear least-squares fit of the helix model **r**(*t*) to the trajectory data, minimising the sum of squared distances between the modelled and simulated positions. The optimisation is performed with a Levenberg–Marquardt algorithm (`LsqFit.jl`), optionally after a temporal smoothing of the trajectory and, where needed, extension over several periods to improve phase estimation.

## Appendix A: Numerical method validation

First, we repeat the convergence tests of [29] for a translating and rotating unit sphere at viscosity *µ* = 1 and *ϵ* = 0.02, where the relative error values show the same convergence as the MATLAB implementation (see Tab. I and Tab. II)

To validate the method for non-spherical geometries, we compute the grand resistance matrix for prolate spheroids and compare against the exact analytical resistance functions of Oberbeck [69]. For a prolate spheroid with semi-major axis *c* and semi-minor axes *a*, the translational and rotational resistance coefficients are

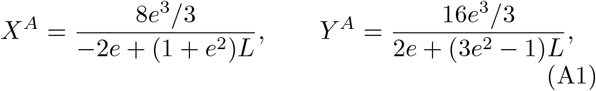

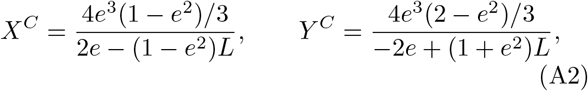

where 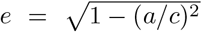 is the eccentricity and *L* = ln((1 + *e*)*/*(1 − *e*)), such that the axial and transverse translational drag forces are *F* = 6*πµc X*^*A*^*U* and *F* = 6*πµc Y* ^*A*^*U* respectively, and the axial and transverse rotational drag torques are *T* = 8*πµc*^3^ *X*^*C*^Ω and *T* = 8*πµc*^3^ *Y* ^*C*^Ω. Surfaces were discretised using a Fibonacci point cloud scaled to the ellipsoid geometry. Results for aspect ratios 2:1 and 5:1 are shown in Tab. III; the maximum error was 0.2%.

**TABLE I.** Relative error in total force (equal to 6*π*) on a translating unit sphere with *µ* = 1, for surface discretisations *N*_*s*_ and quadrature orders *Q*_*s*_.

| $N_s$ | $Q_s=864$ | 3456 | 13824 | 55296 | 221184 |
| --- | --- | --- | --- | --- | --- |
| 54 | 0.758% | 0.514% | 0.515% | 0.514% | 0.514% |
| 216 | 0.901% | 0.541% | 0.515% | 0.515% | 0.515% |
| 864 | 4.676% | 0.579% | 0.516% | 0.515% | 0.515% |
| 3456 | 0.614% | 0.271% | 0.517% | 0.515% | 0.515% |

**TABLE II.** Relative error in total torque (equal to 8*π*) on a rotating unit sphere, with *µ* = 1 for surface discretisations *N*_*s*_ and quadrature orders *Q*_*s*_.

| $N_s$ | $Q_s=864$ | 3456 | 13824 | 55296 | 221184 |
| --- | --- | --- | --- | --- | --- |
| 54 | 1.657% | 1.220% | 1.219% | 1.211% | 1.211% |
| 216 | 2.395% | 1.528% | 1.487% | 1.487% | 1.486% |
| 864 | 8.132% | 1.670% | 1.521% | 1.519% | 1.519% |
| 3456 | 1.712% | 0.055% | 1.528% | 1.522% | 1.522% |

**TABLE III.** Comparison of numerical and analytical resistance matrix entries for prolate spheroids of aspect ratio 2:1:1 and 5:1:1, with *µ* = 1 and semi-major axis *c* equal to the aspect ratio. Analytical values are from the exact Oberbeck solution [69]. Numerical results use 531 force points and 50539 quadrature points with *ϵ* = 10^*−*5^.

| | $6\pi\mu c \cdot X^A$ | $6\pi\mu c \cdot Y^A$ | $8\pi\mu c^3 \cdot X^C$ | $8\pi\mu c^3 \cdot Y^C$ |
| --- | --- | --- | --- | --- |
| Analytical (2:1:1) | 22.694 | 25.992 | 40.548 | 75.646 |
| Numerical (2:1:1) | 22.702 | 26.006 | 40.600 | 75.726 |
| Error | 0.04% | 0.06% | 0.13% | 0.11% |
| Analytical (5:1:1) | 33.643 | 44.692 | 88.729 | 583.143 |
| Numerical (5:1:1) | 33.660 | 44.729 | 88.910 | 584.149 |
| Error | 0.05% | 0.08% | 0.20% | 0.17% |

We also calculated the total force on a straight flagellum of length *L* = 1, radius *a* = 0.01 translating perpendicular or parallel to its tangent with unit speed (keeping viscosity *µ* = 1), and compared these with the leading-order slender-body drag coefficients [70, 71]

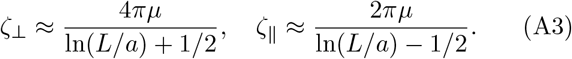

The results are shown in Tab. IV and Tab. V. We took the regularisation parameter *ϵ* ≈ 1.118 *a*, which provides a reasonable match (*<* 1% error) to both coefficients simultaneously.

**TABLE IV.** Relative error in total force for a straight filament translating perpendicular to its axis.

| $N_f$ | $Q_f=128$ | 256 | 512 | 1024 |
| --- | --- | --- | --- | --- |
| 32 | 0.810% | 1.061% | 1.187% | 1.250% |
| 64 | 0.264% | 0.500% | 0.617% | 0.675% |
| 128 | - | 0.018% | 0.119% | 0.169% |
| 256 | - | - | 0.342% | 0.301% |

**TABLE V.** Relative error in total force for a straight filament translating parallel to its axis.

| $N_f$ | $Q_f=128$ | 256 | 512 | 1024 |
| --- | --- | --- | --- | --- |
| 32 | 0.886% | 1.108% | 1.218% | 1.269% |
| 64 | 0.128% | 0.335% | 0.437% | 0.487% |
| 128 | - | 0.282% | 0.194% | 0.149% |
| 256 | - | - | 0.722% | 0.684% |

We test of convergence of the three example organisms using behaviour metrics considered to have converged when the relative change in the quantity upon doubling the discretisation is *O*(10^−4^) (thus an error of *O*(1) only accrues after 10^4^ beat cycles). For the trajectory measurements of the *Chlamydomonas* example (Sec. III A) and the morphological transition (Sec. III C) we use the displacement over the beat cycle, and for the filter feeder (Sec. III B) we use the feeding flux and dissipated power. The discretisations used can be found in Tab. VI.

**TABLE VI.** Converged values for the quadrature and force point discretisations.

| Section | $N_b$ | $Q_b$ | $N_f$ | $Q_f$ |
| --- | --- | --- | --- | --- |
| III A | 237 | 999 | 87 | 261 |
| III B | 3131 | 25048 | 97 | 388 |
| III C | 237 | 3996 | 94 | 522 |

To check convergence at a finer level of accuracy, we can exploit the two discretisations. We compute the velocity field using the calculated force distribution (for a `ResistanceProblem` or `SwimmingProblem`). This force distribution is constant over each patch corresponding to a force point. Since we only impose the velocity at force points, a useful diagnostic is the predicted velocity on the finer quadrature grid. We compute the residual of the prescribed velocity at the quadrature points (not used in the solution) with the predicted velocity given by the fluid velocity at those points, relative to the magnitude of the prescribed velocity. Our chosen discretisations respect that the median value of this residual is *O*(10^−3^) and the 95th quartile *O*(10^−2^).

